# Evidence for a reversal of the neural information flow between object perception and object reconstruction from memory

**DOI:** 10.1101/300913

**Authors:** Juan Linde-Domingo, Matthias S. Treder, Casper Kerren, Maria Wimber

## Abstract

Remembering is a reconstructive process. Surprisingly little is known about how the reconstruction of a memory unfolds in time in the human brain. We used reaction times and EEG time-series decoding to test the hypothesis that the information flow is reversed when an event is reconstructed from memory, compared to when the same event is initially being perceived. Across three experiments, we found highly consistent evidence supporting such a reversed stream. When seeing an object, low-level perceptual features were discriminated faster behaviourally, and could be decoded from brain activity earlier, than high-level conceptual features. This pattern reversed during associative memory recall, with reaction times and brain activity patterns now indicating that conceptual information was reconstructed more rapidly than perceptual details. Our findings support a neurobiologically plausible model of human memory, suggesting that memory retrieval is a hierarchical, multi-layered process that prioritizes semantically meaningful information over perceptual detail.

## 1. Introduction

When Rocky Balboa goes back to his old gym in the film Rocky V, the boxing ring and the feeling of the dusted gloves in his hands trigger a flood of vivid images from the past. Like in many other movies featuring such mnemonic flashbacks, the main character seems capable of remembering what the room looked like years ago, who was there at the time, and even an emotional conversation with his old friend and coach Michael. Perceptual details like colours, however, are initially missing in the scene, like in a faded photograph, and only gradually saturate over time. This common way to depict memories in pop culture nicely illustrates that the memories we bring back to mind are likely not unitary constructs, and also not veridical copies of past events. Instead, they suggest that remembering is a reconstructive process that might prioritize more meaningful components of an event over other more shallow aspects (Schacter, 2012; Schacter, Guerin, & St Jacques, 2011). We here report three experiments that shed light onto the temporal information flow during memory retrieval. Once a reminder has elicited a stored memory trace, are the different features of this memory reconstructed in a systematic, hierarchical way?

Considering our vast knowledge about the information processing hierarchy during visual perception, surprisingly little is known about the time course of memory recall. In the object recognition literature, it is generally agreed that the presentation of an external stimulus initiates a processing cascade that starts with low-level perceptual features in early visual areas, and progresses to increasingly higher levels of semantic integration and abstraction along the inferior temporal cortex (Carlson, Tovar, Alink, & Kriegeskorte, 2013; Cichy, Pantazis, & Oliva, 2014; Clarke & Tyler, 2015; Lehky & Tanaka, 2016; Martin, Douglas, Newsome, Man, & Barense, 2018; Serre, Oliva, & Poggio, 2007). However, mental representations can also be re-created from memory, without much external stimulation: retrieving a scene from the movie Rocky V will elicit semantic knowledge about the film (e.g. that the actor is called Sylvester Stallone), but also mental images that can include fairly low-level details (e.g. whether the scene was in colour or in grey scale). How the brain manages to bring back each of these features when reconstructing an event from memory remains an open question. The present series of experiments tested our central working hypothesis that the stream of information processing is reversed during memory reconstruction compared with the perception of an external stimulus.

Over the last years, multivariate neuroimaging methods have made it possible to isolate brain activity patterns that carry information about externally presented stimuli, but also about internally generated mnemonic representations. Importantly, it has been shown that the neural trace that an event produces during its initial encoding is reinstated in brain activity during its later retrieval (Chen et al., 2017; Johnson, McDuff, Rugg, & Norman, 2009; Kuhl, Rissman, Chun, & Wagner, 2011; Michelmann, Bowman, & Hanslmayr, 2016; Staresina, Henson, Kriegeskorte, & Alink, 2012; Wimber, Alink, Charest, Kriegeskorte, & Anderson, 2015). Most of these studies focused on the reactivation of abstract information, including a picture’s category (Kuhl et al., 2011; Staresina et al., 2012; Wimber et al., 2015) or the task context in which it was encoded (Johnson et al., 2009). Apart from these higher-level features, evidence also exists for the reactivation of low-level perceptual details in early visual areas (Bosch, Jehee, Fernandez, & Doeller, 2014; Waldhauser, Braun, & Hanslmayr, 2016). Moreover, a growing literature using electrophysiological methods has begun to shed light onto the timing of such reinstatement, typically demonstrating neural reactivation within the first second after a reminder is presented (Jafarpour, Fuentemilla, Horner, Penny, & Duzel, 2014; Michelmann et al., 2016; Sols, DuBrow, Davachi, & Fuentemilla, 2017; Staudigl et al., 2012), and sometimes very rapidly (Waldhauser et al., 2016; Wimber, Maaß, Staudigl, Richardson-Klavehn, & Hanslmayr, 2012). However, because all existing studies only focused on a single feature of a memory representation (e.g., its semantic category), the fundamental question whether memory reconstruction follows a hierarchical information processing stream, similar to perception, has not been investigated.

We hypothesize that such a processing hierarchy does exist, and that the information flow is reversed during memory reconstruction compared with perception. That is, based on the widely accepted idea that memory reconstruction depends on back-projections from the hippocampus to neo-cortex (Moscovitch, 2008), we expect that those areas that are anatomically closer to the hippocampus (i.e. high-level conceptual processing areas along the inferior temporal cortex) should be involved in the reactivation cascade faster that areas that are relatively remote (i.e., low-level perceptual processing areas in earlier visual cortices). Therefore, we assume that once a reminder has initiated the reactivation of an associated event, higher-level abstract information will be reconstructed before lower-level perceptual information, producing an inverse temporal order of processing compared with perception.

We tested this reverse reconstruction hypothesis in a series of two behavioural and one EEG experiment (see Fig. 1b, c, and Fig. 3a). All experiments used a simple associative memory paradigm where participants learn a series of arbitrary associations between word cues and everyday objects, and are later cued with the word to recall the object. In order to test for a processing hierarchy, it is important to independently manipulate the perceptual and conceptual contents of these objects. Therefore, objects varied along two orthogonal dimensions: one perceptual dimension, where the object can be either presented as a photograph or a line drawing; and a semantic dimension where the object represents an animate or inanimate entity (Fig. 1a). The two behavioural experiments measure reaction times while participants make perceptual or semantic category judgments for objects that are either visually presented on the screen, or reconstructed from memory. The EEG experiment uses a similar associative recall paradigm together with time-series decoding techniques (Carlson et al., 2013; Cichy et al., 2014; Kurth-Nelson, Barnes, Sejdinovic, Dolan, & Dayan, 2015), allowing us to track at which exact moment in time perceptual and semantic components of the same object are reactivated, and to create a temporal map of semantic and perceptual features during perception and memory reconstruction (Fig. 3b and c). Our behavioural and electrophysiological findings consistently support the idea that memory reconstruction is not an all-or-none process, but rather progresses on each single trial from higher-level semantic features to lower-level perceptual details.

**Figure 1.**
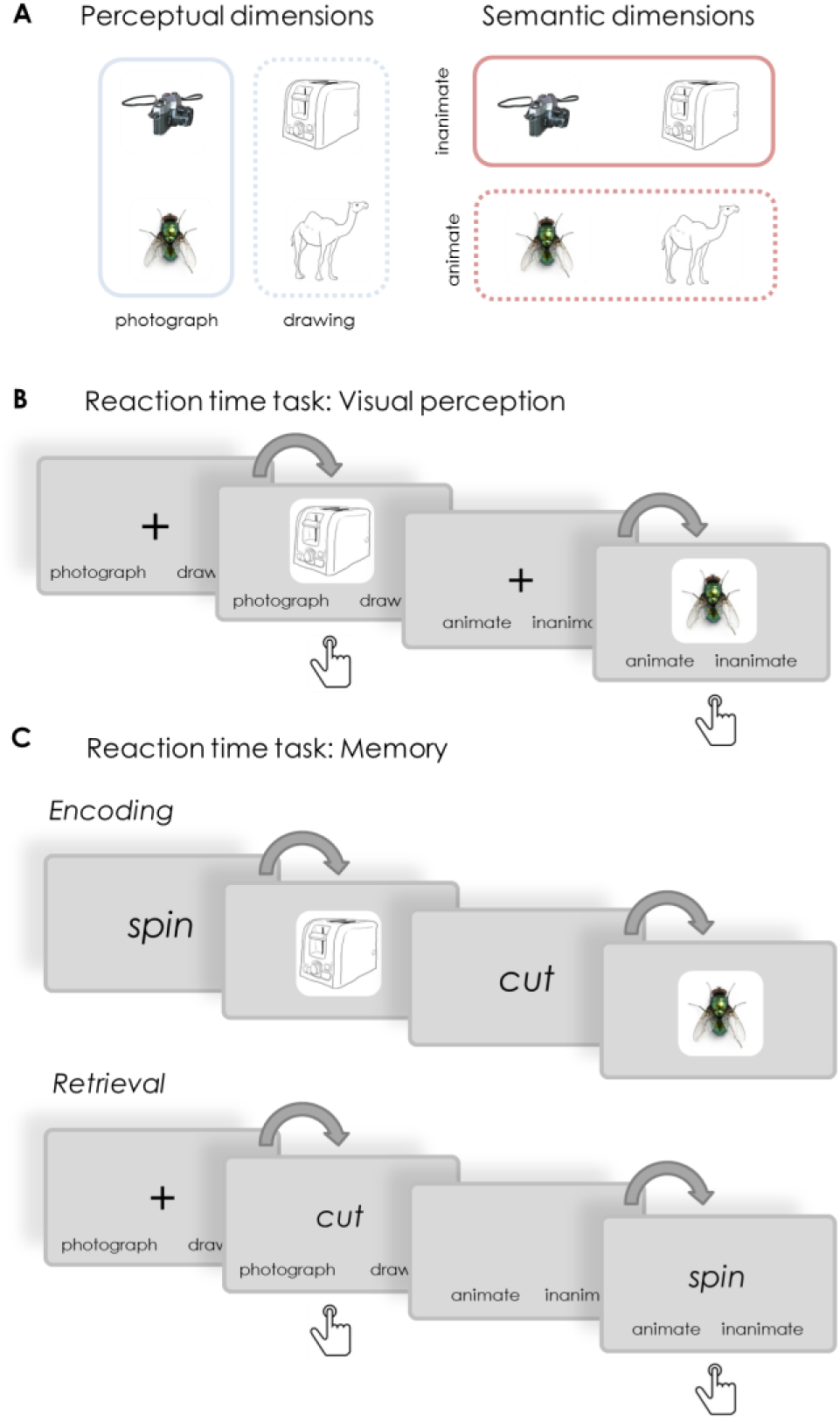
Stimuli and design of the behavioural experiments. (a) Illustration of the orthogonal design of the stimulus set. In all experiments, objects (a total of 128) varied along two dimensions: a perceptual dimension where objects could be presented as a photograph or as a line drawing; and a semantic dimension where objects could belong to the animate or inanimate category. (b) In the visual reaction time task, participants were prompted on each trial to categorize the upcoming object as fast as possible, either according to its perceptual category (photograph vs. line drawing) or its semantic category (animate vs. inanimate). (c) During the encoding phase of a memory reaction time task, participants were asked to create word-object associations (a total of 8 per block). Reaction times were then measured during the retrieval phase, where subjects were presented with a reminder word, and asked to recall and categorize the associated object according to its perceptual (photograph vs. line drawing) or semantic (animate vs. inanimate) features. Button press symbols indicate at which moment in a trial RTs were collected.

## 2. Results

### 2.1. Behavioural experiments

Our two behavioural experiments used reaction times (RTs) to test our central hypothesis that the information processing hierarchy reverses between the visual perception of an object, and its reconstruction from memory. We assumed that the time required to answer a question about low-level perceptual (photograph vs. drawing) compared to high-level semantic (animate vs. inanimate) features of an item would reflect the speed at which the brain gains access to these types of information. If so, we expected that reaction time patterns would reverse depending on whether the object is visually presented or reconstructed from memory: during visual perception, RTs should be faster for perceptual compared with semantic questions to mirror the forward processing hierarchy, while during retrieval RTs should be faster for semantic compared with perceptual questions if there is a reversal of that hierarchy.

Both experiments used a 2 × 2 mixed design (Fig. 1b and c), where all participants answered perceptual and semantic questions (factor question type, within-subjects) about the objects. Importantly, one group of participants was visually presented with the objects while answering these questions, whereas the other group recalled the same objects from memory (factor task, between-subjects). The main difference between the two experiments was that in Experiment 1, both types of features were probed for a given object, and that in Experiment 2, object were presented on background scenes (not of interest for the present purpose; see Methods section for details). Overall accuracy in both experiments was near ceiling for the visual reaction time task (Experiment 1: M = 96.88%; SD = 2.40%; Experiment 2: M = 97.19%, SD = 2.99%), and high for the memory reaction time task (Experiment 1: 83.15%; SD = 0.92; Experiment 2: M = 66.23%, SD = 15.35). Only correct trials were used for all further RT analyses.

#### 2.1.2. Reaction times show the expected reversal in Experiments 1 and 2

To directly test for a reversal of the reaction time pattern between visual perception and memory reconstruction, we performed an analysis of variance comparing the RTs to perceptual and semantic questions during visual object presentation and during the cued-recall task. As predicted, we found a significant interaction between task (visual vs. memory group) and question type (i.e. perceptual vs. semantic) in Experiment 1 (*F*_1,42_ = 11.142, *P* = .002) and in Experiment 2 (*F*_1,46_ = 10.876, *P* = .002). There was no main effect of question type (Experiment 1: *F*_1,42_ = 3.816, *P* = .057; Experiment 2: *F*_1,46_ = 3.184, *P* = .081), suggesting that participants were not generally faster or slower at answering one type of question compared to the other (Fig. 2a and b).

**Figure 2.**
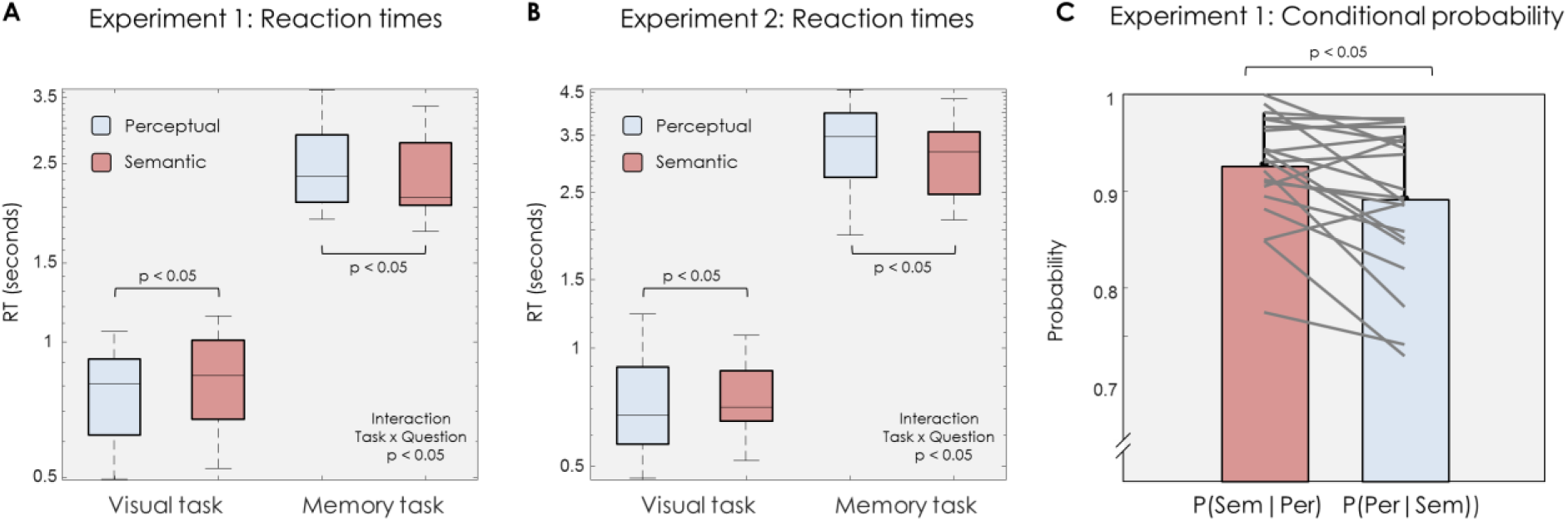
Behavioural results. (a) Box plots representing reaction times in Experiment 1 for perceptual (blue) and semantic questions (pink) during object presentation (visual task, left) and object recall (memory task, right). A significant interaction was found in a 2×2 ANOVA comparing RTs for perceptual and semantic questions when an object was physically presented on the screen (visual task) or cued by a reminder (memory task). (b) Box plots representing reaction times in Experiment 2 for perceptual and semantic questions during in the visual and memory groups, replicating the results from Experiment 1. For illustrative purposes the Y-axis in (a) and (b) is logarithmically scaled. (c) Conditional probability results in Experiment 1. The conditional probability of remembering the correct semantic information given the perceptual question was answered correctly for the same object, P(sem/per) was significantly higher than the conditional probability of answering the perceptual question correctly given a correct semantic answer, P(per/sem). Each line represents the trend for one participant. In all three panels, errors bars represent standard error of the mean. The line in the middle of each box represents the median, and the tops and bottoms of the boxes the 25th and 75th percentiles of the samples, respectively. Whiskers are drawn from the interquartile ranges to the furthest minimum (bottom) and maximum (top) values.

Post-hoc RT analyses were then performed for each task to confirm that this interaction was produced by differences in the expected direction. In Experiment 1, participants in the visual perception group were significantly faster when responding to perceptual (M = 795ms; SD = 235ms) compared to semantic (M = 842ms, SD = 185ms) questions (*t*_22_ = 3.68, *P* = .001). Importantly, these differences reversed in the memory retrieval group, where RTs to semantic questions (2334ms; SD = 534 sec) were now significantly faster than RTs to perceptual questions (M = 2502ms; SD = 561; *t*_20_ = 2.35, *P* = .029). This pattern was fully replicated in Experiment 2, where again the visual RT group answered perceptual questions (M = 733ms; SD = 211ms) significantly faster than semantic questions (M = 797ms, SD = 235; *t*_23_ = 2.46, *P* = .022), whereas the memory group was significantly faster at responding to semantic questions (M = 3133ms, SD = 660ms)) compared with perceptual questions (M = 3348ms, SD = 754; *t*_23_ = 2.67, *P* = .014).

Since reaction times are not necessarily normally distributed, we also wanted to confirm the results using a Wilcoxon signed rank test. The significant RT differences between perceptual and conceptual questions were also present using this non-parametric statistic in the visual perception group (Experiment 1: *z* = 3.16, *P* = .002; Experiment 2: *z* = 2.57, *P* = .010) and in the memory group (Experiment 1: *z* = 2.48, *P* = .013; Experiment 2: *z* = 2.42, *P* = .015). Reaction time analyses thus support our central hypothesis that the speed of information processing for different object features reverses between perception and memory, and this pattern fully replicated between Experiments 1 and 2.

#### 2.1.3. Accuracy results support a reversal between perception and memory, and suggest a directional dependency in the processing hierarchy

Next we investigated whether a similar pattern was, at least qualitatively, also present in terms of accuracy. We found a significant interaction between task (visual vs. memory group) and question type (i.e. perceptual vs. semantic question) in both experiments (Experiment 2: *F*_1,42_ = 14.467, *P* = .001; Experiment 2: *F*_1,46_ = 9.698, *P* = .003). Post-hoc accuracy analyses in Experiment 1 revealed that in the visual reaction task participants were more accurate at answering perceptual questions (M = 97.42%; SD = 2.68%) compared to semantic ones (M = 96.33%; SD = 1.99%). This difference was not significant (*t*_22_ = 2.03, *P* = .055), most likely because accuracy during perception was close to ceiling. Accuracy in the memory task showed that, in line with a reversed processing stream, participants had significantly better accuracy for semantic questions compared with perceptual questions (M = 85.83%; SD = 7.57%; vs. 82.63%; SD = 8.79%, respectively; *t*_20_ = 3.12, *P* = .005). Experiment 2 replicated the same accuracy profile, with participants in the visual group showing a significantly higher accuracy for perceptual questions (M = 97.97%; SD = 2.77%) compared to semantic questions (M = 96.41%; SD = 3.07%; *t*_23_ = 2.14, *P* = .042)). The reverse pattern was present in the memory reaction time task, where an accuracy benefit was found for semantic questions compared to perceptual ones (69.57%; SD = 15.17%; vs. 62.89%; SD = 15.09%, respectively; *t*_23_ = 2.63, *P* = .015). Accuracy profiles thus generally corroborated our reaction time results, again suggesting that semantic information is more easily accessed during retrieval than perceptual information.

The accuracy data from Experiment 1 also allowed us to address an interesting question regarding the dependency of perceptual and conceptual processing stages. Across the retrieval phase of this experiment, both types of questions were asked for each given object, and we were thus able to test to what degree performance on the semantic and perceptual questions was stochastically dependent. Our reasoning was that if the reconstruction of semantic and perceptual aspects from memory was a hierarchical process where access to a later stage depended on having completed the previous stage(s), as predicted by a reversed stream, then the ability to retrieve perceptual details would depend on having accurately retrieved semantic details, but not vice versa. In other words, if the retrieval of semantic information was the first stage in a hierarchical stream, it would not depend much on any other stages. If on the other hand the retrieval of perceptual information is indeed a very late stage in the hierarchy, success at this stage should be considerably influenced by success at earlier (semantic) stages. In line with this reasoning, we found that P(sem/per) – the conditional probability of remembering the correct semantic information given the perceptual question was answered correctly for the same word-picture association (M = 91.61%; SD = 6.98%) – was significantly higher (*t*_20_ = 3.08, *P* = .006) than P(per/sem) – the conditional probability of answering the perceptual question correctly given a correct semantic answer (M = 88.28%; SD = 8.34%)(Fig. 2c). For reasons of completeness, we carried out the same conditional probability analysis in the visual task. In this group, the opposite trend was present, with P(per/sem) (M = 97.30%; SD = 2.82 %) being numerically higher than P(sem/per) (M = 96.21%; SD = 2.09%). However, this difference was not statistically robust (*t*_22_ = 2.04, *P* = .054), most likely due to ceiling effects.

Altogether, the findings from our two behavioural experiments provide support for our main hypothesis that during retrieval of a complex visual representation, the temporal order in which perceptual and semantic features are processed reverses between perception (feed-forward) and memory retrieval (feed-backward). The results suggest that reaction times can be used as a proxy to probe neural processing speed, as argued in previous studies (Ritchie, Tovar, & Carlson, 2015). In the next sections, we report the findings from an EEG study that more directly taps into the neural processes that we believe are producing the behavioural pattern.

### 2.2. EEG experiment

While it is reasonable to assume that reaction times tap into the neural processing speed for a given feature, based on previous literature (Ritchie et al., 2015), we also wanted to obtain a more direct signature of feature activation from human brain activity. We therefore used multivariate pattern analysis applied to electrophysiological (EEG) recordings, with the goal to pinpoint when in time, on an individual trial, the perceptual and semantic features of an object could be decoded from brain activity. We expected to find the maximum decodability of perceptual information before semantic information when an object was visually presented on the screen, and expected the order of these peaks to reverse when the object was recalled from memory. The design closely followed the behavioural experiments, with the important difference that all factors were manipulated within subjects, such that each participant carried out a visual encoding phase that served to probe visual (forward) processing, and a subsequent recall phase used to probe mnemonic (backward) processing (Fig. 3). The trial timing was optimised for obtaining a clean signal during object presentation and object recall, rather than for measuring reaction times. We therefore presented the perceptual and semantic questions only during the recall phase in order to probe memory accuracy, and questions were presented at the end of each recall trial, such that they would not bias processing towards perceptual or semantic features of the object.

**Figure 3.**
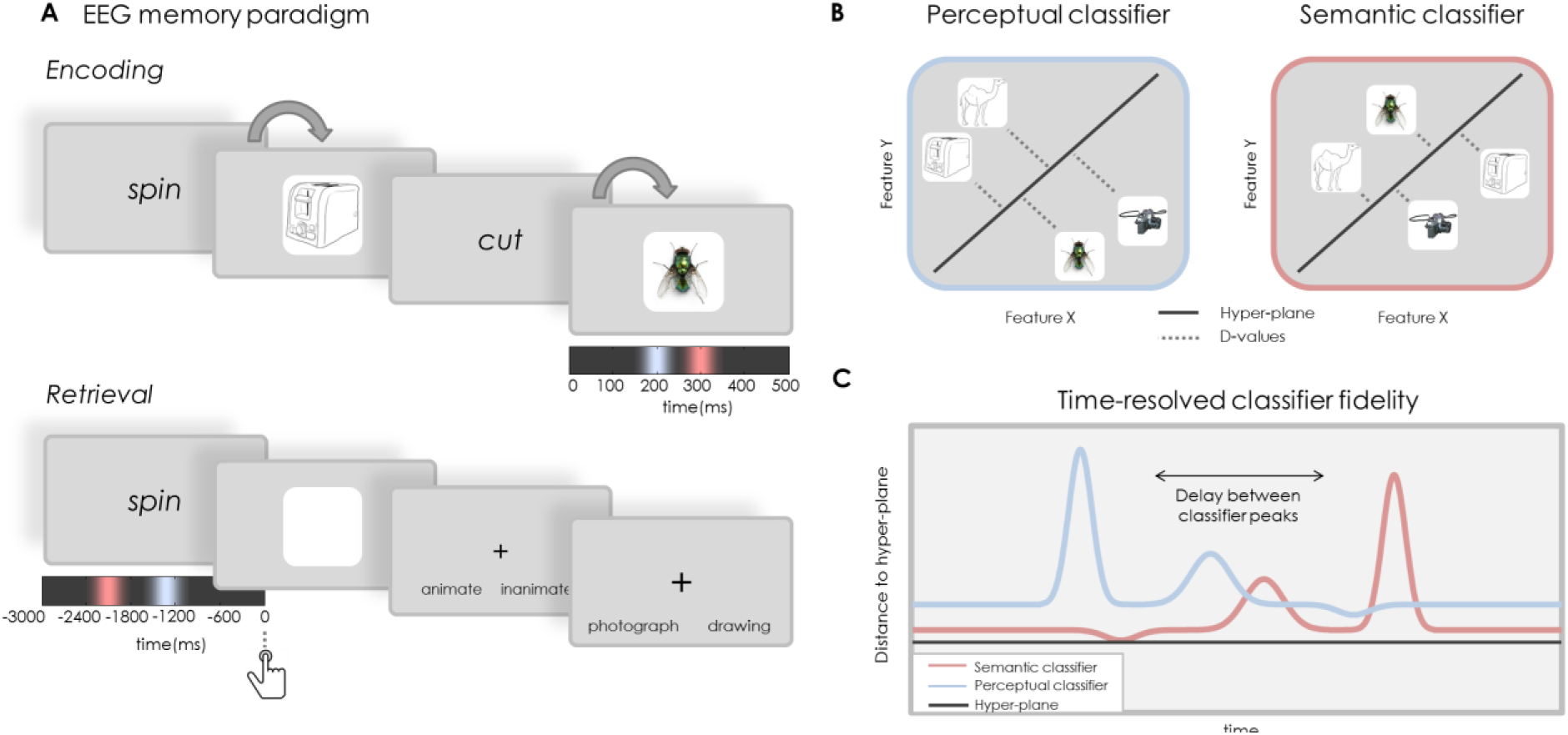
Design for EEG experiment and time resolved multivariate decoding. In the EEG experiment participants were asked to create word-object associations (panel A), and to later reconstruct the object as vividly as possible when cued with the word, and to indicate with a button press when they had a vivid image back in mind. EEG was recorded during learning and recall, with the aim to perform time-series decoding analyses that can detect at which moment, within a single trial, a classifier is most likely to categorise perceptual and semantic features correctly. Coloured time lines under object and cue time windows represent our reversal hypothesis regarding the temporal order of maximum semantic (pink) and perceptual (blue) classification during the perception (encoding) and retrieval of an object. All EEG analyses were aligned to the object onset during encoding, and to the button press during retrieval. (b) Decoding analyses were performed independently per participant at each time point. For each given time point during a trial, two linear discriminant analysis (LDA) based classifiers were trained on the EEG signal: one perceptual classifier discriminating photographs from line drawings, and one semantic classifier discriminating animate from inanimate objects. Classifiers were tested using a leave-one-out procedure, which allowed us to obtain a time series of confidence values (d-values, reflecting the distance from the separation hyperplane) for each single trial. (c) Our main interest was to compare the time points of maximal fidelity of the perceptual (blue) and semantic classifiers (pink) on each trial, to test the hypothesis that the perceptual maximum (blue) precedes the semantic one (pink) during perception (encoding, panel A), and importantly that this order is reversed during memory recall (panel B).

#### 2.2.1 Accuracy in the EEG study replicates the response pattern found in the behavioural experiments

In the retrieval phase of the EEG experiment, subjects were again cued with a word and asked to retrieve the associated object. On average participants subjectively declared to retrieve the object on 93.6% of the trials (SD = 5.89%), with an average reaction time of 3046ms (SD = 830ms; minimum = 1369ms; maximum = 5124ms) to make this response. We then asked two objective questions at the end of each trial, one perceptual and one semantic, which participants answered with an overall mean accuracy of 86.37% (SD = 6.6). Mirroring our behavioural experiments, hit rates for answering the semantic question were 87.65% (SD = 6.57%), significantly higher (*t*_20_= 5.16, *P* = .001) than the accuracy for the perceptual question (M = 85.08%; SD = 6.53%). Note that the EEG task was not designed to measure reaction times, and participants were instructed to prioritize accuracy over speed.

#### 2.2.2 Single-trial classifier fidelity suggests a reversal of information processing between perception and memory recall

In order to determine the temporal trajectory of feature processing on a single trial level, we carried out a series of time resolved decoding analyses. Linear discriminant analysis (LDA, see Method section) was used to classify perceptual (photograph vs. line drawing) and semantic (animate vs. inanimate) features of an object based on the EEG topography at a given time point, either during object presentation (encoding) or during object retrieval from memory (cued recall).

Our first aim was to confirm that there was a forward stream during perceptual object processing. Two separate classifiers were therefore trained and tested during encoding to classify the perceptual category (photograph vs. line drawing) and the semantic category (animate vs. inanimate) of the to-be-encoded object, respectively, in each trial and time point per participant (see Fig. 3). For these analyses, decoding was performed in separate time windows starting 100ms before stimulus onset and up until 500ms post-stimulus. Our main interest was to determine the specific moment in each trial at which the two classifiers showed the highest fidelity in determining the correct perceptual and semantic categories (Fig. 3b and c). For the encoding data, we thus identified the highest *d* value peak per trial within 500ms of stimulus onset (see Methods section). This approach allowed us to compare, within each encoding trial, whether the classification peak for perceptual features occurred earlier than the classification peak for semantic features.

Comparing all single trial *d* value peaks from encoding (Fig. 4a), we found a significant difference (*z* = 1.87, *P* = 0.03) between the timing of perceptual and semantic peaks using a one-tailed Wilcoxon signed rank test, suggesting that confidence peaks for perceptual classification occurred before those for semantic classification. The obtained Z score was compared against a bootstrapped data set (see Methods section) to estimate the likelihood of obtaining a distance between peaks of the same or larger size from a distribution with randomly shuffled category labels, using the same EEG epochs and the same time window. The observed difference score (*z* = 1.87) exceeded the 95^th^ percentile (*z* = 1.64) of the random distribution. This result from the encoding phase of the experiment thus confirms previous studies showing that low-level features are processed before high-level features during visual perception (Carlson et al., 2013; Cichy et al., 2014; Clarke & Tyler, 2015; Lehky & Tanaka, 2016; Serre et al., 2007).

**Figure 4.**
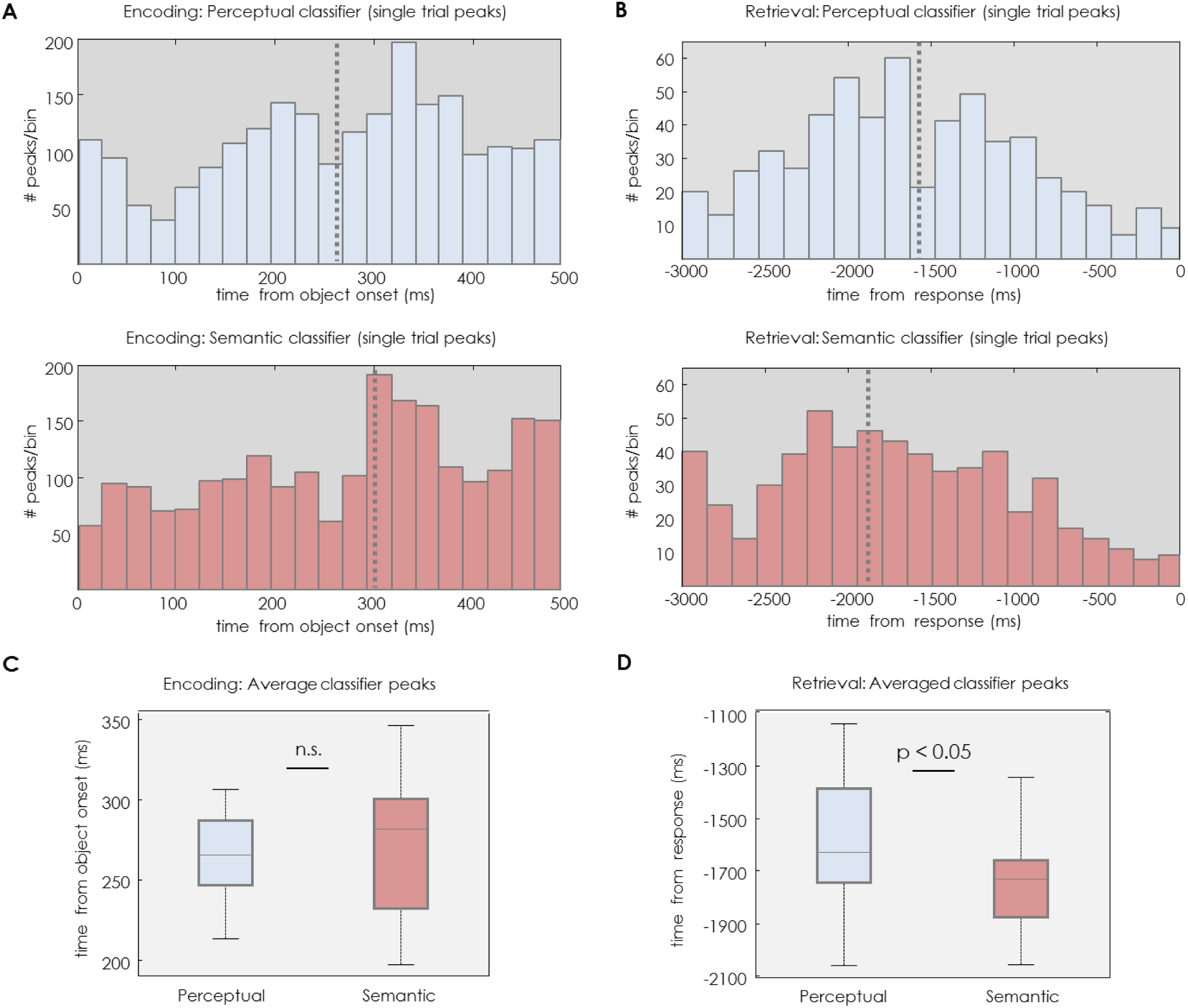
EEG multivariate analysis results. Classifier fidelity in terms of single-trial ***d*** value peak distributions (dashed lines represent the median) during object encoding (a) and object retrieval (b), shown separately for classifying perceptual (blue) and semantic (pink) classes. A significant difference between the two peak distributions was found during object encoding (*P* = .015), indicating a bias towards earlier occurrence of perceptual (blue) compared with semantic (pink) peaks. During object retrieval, a significant difference between the distributions was found (*P* = .006) in the opposite direction relative to encoding, with semantic peaks now occurring earlier than perceptual peaks. Box plots represent group peak distribution of *d* values for perceptual and semantic categories during encoding (c) and retrieval (d) after averaging peaks within participants. A significant interaction (P = .048) was found between task (encoding or retrieval) and type of feature (i.e. perceptual or semantic). n.s. indicates a non–significant T-value in posthoc tests. All box plots elements represent the same metrics as in Figure 2.

Importantly, following the same procedure, we next analysed the differences between the perceptual and semantic classifier peaks during memory reactivation, to test whether the order reversed during retrieval compared with encoding. The single-trial approach made sure that the relative temporal order of perceptual and semantic peaks within a trial would be preserved even if the retrieval process was set off with a varying delay across trials. To further minimize variance between the retrieval trials, we aligned all trials relative to the button press, i.e. the moment when participants declared that they had retrieved the associated object from memory. The time window used in this analysis covered 3sec prior to participants’ response and, based on behavioural reaction times, only trials with a RT ≥ 3 sec were included. Using a one-tailed Wilcoxon signed rank test, a significant difference (*z* = 2.53, *P* = .006) was found when we compared *d* value peak distributions of perceptual with those of semantic classification obtained from all single trials and participants (Fig. 4b). The obtained Z score was again higher than the 95^th^ percentile (*z* = 1.59) of the random distribution of a bootstrapped data set (see Methods section) using the same EEG signal and time window. Critically, the one-tailed test in this case confirms our central hypothesis that during memory retrieval, semantic information can be classified in brain activity significantly earlier than perceptual information, suggesting a reversal of information flow relative to perception.

The last classification analysis was aimed at confirming the results obtained from the previous single-trial, fixed-effects analyses using a random-effects approach. We calculated the average *d* value peak latency for perceptual and semantic classification in each participant, and performed a 2×2 ANOVA with stage (encoding vs. retrieval) and type of feature (perceptual vs. semantic) as within-subject factors. Confirming our main hypothesis, this analysis revealed a significant interaction (*F*_1,20_ = 4.63, *P* = .044) between stage and the type of feature. We further found a main effect of type of feature (*F*_1,20_ = 4.80, *P* = .04). Post-hoc T-tests showed no significant difference (*t*_20_= 0.67, *P* = .253) between the average perceptual and semantic *d* value peaks during encoding (Fig. 4c). However, during retrieval, we found that the semantic classifier systematically (*t*_20_= 2.20, *P* = .020, one-tailed) peaked earlier than the perceptual classifier (Fig. 4d). These findings indicate that even though a single-trial comparison of classifier fidelity is more sensitive to the temporal dynamics of feature processing, the same pattern is also present in the average classification values.

Overall, the results again confirm our hypothesis that the information processing hierarchy reverses between perception (encoding) and recall, and that memory recall prioritizes semantic over perceptual information.

#### 2.2.3 Univariate ERP results are consistent with the reverse processing hypothesis

In a final step, we also sought to corroborate our findings by more conventional event-related potential (ERP) analyses. If the differences in neural activity between perceptual (photograph vs. line drawing) and semantic (animate vs. inanimate) categories, as picked up by the LDA classifier, were produced by a signal that is relatively stable across trials and participants, we expected to see these signal differences in the average ERP time courses across participants. A comparison of the ERP peaks during encoding and retrieval would then reveal the same perception-to-memory reversal as found in our multivariate analyses.

Firstly, a series of cluster-based permutation tests (see Methods section) was performed during object presentation to test for ERP differences between perceptual and semantic categories. Contrasting objects from the two different perceptual categories (photographs and line drawings), we obtained a significant positive cluster (*P*_corr_ = .008) between 136ms and 232ms after stimulus onset, with a maximum difference based on the sum of T values at 188ms, and located over occipital and central electrodes (see Fig. 5a). Contrasting objects from the different semantic categories (animate and inanimate) revealed a later cluster over frontal and occipital electrodes (*P*_corr_ = .001) from 237ms until 357ms after stimulus presentation, with a maximum difference at 306ms (see Fig. 5a). The peak semantic ERP difference for encoding thus occurred ^~^120ms after the peak perceptual difference, consistent with the existing ERP literature (Fabiani, M., Gratton, G., & Federmeier, 2007).

**Figure 5.**
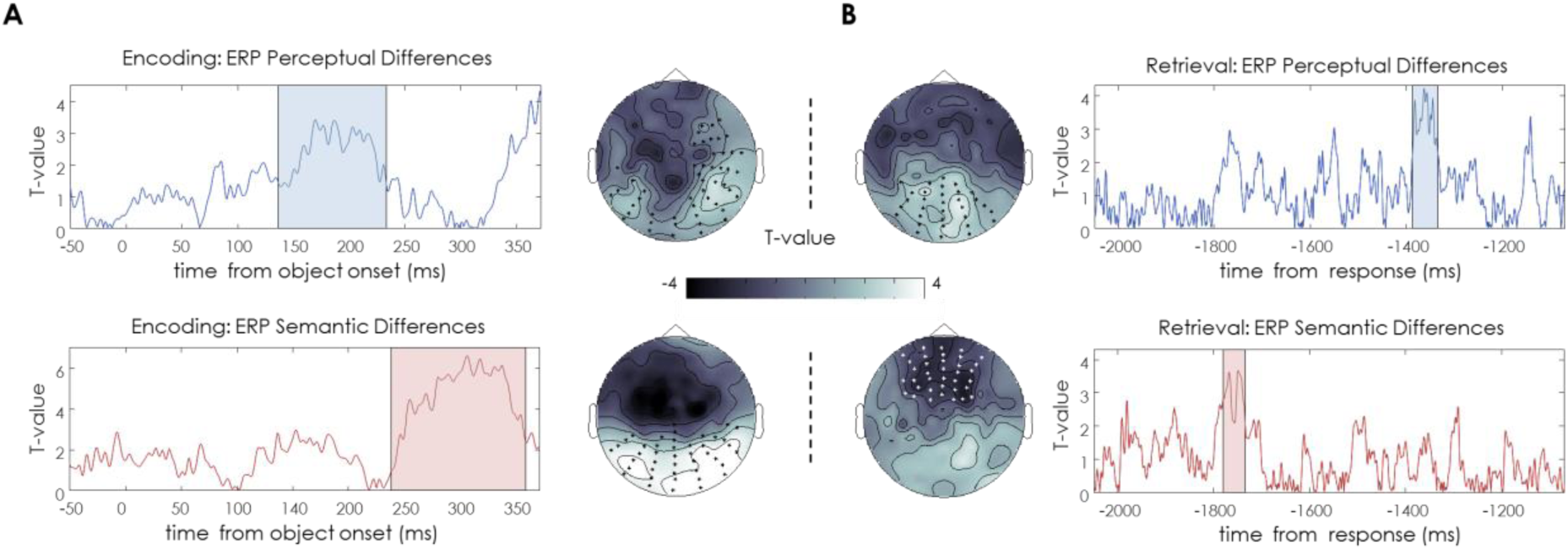
Univariate analysis results. (a) Left panels represent ERP group differences (T-values) across time in those electrodes that formed a significant cluster during object presentation, locked to the onset of the stimulus. Top left panel shows the contrast of photographs vs. line drawings, and the bottom left panel differences between animate vs. inanimate objects. Scalp figures next to each contrast illustrate the maximum cluster’s topography, averaged across the significant time-window, with all significant electrodes in a cluster being marked with an asterisk. (b) Right panels show ERP group differences (T-values) over time in those electrodes that are contained in the maximum significant clusters during memory retrieval, time locked to participants’ responses). The top right panel shows the perceptual contrast, and the bottom right panel the semantic contrast. Cluster topographies for each comparison are located next to each panel, and the temporal extent of significant clusters is shaded in colour.

Similar contrasts between perceptual and semantic categories were then carried out during retrieval, aligning trials to the time of the button press. We found a significant perceptual cluster distinguishing the recall of photographs and line drawings over occipital electrodes (*P*_corr_ = .046) between 1390ms and 1336ms before participants’ responses, with a maximum difference based on the sum of T values at 1360ms prior to response time (see Fig. 5b). Comparing ERPs for the different semantic categories, we found a significant cluster distinguishing the recall of animate from inanimate objects over frontal electrodes *(P_corr_* = .032) between 1781ms and 1735ms before object retrieval, with a maximum difference at 1770ms (see Fig. 5b). Therefore, during memory retrieval, the peak semantic ERP difference occurred ~400ms before the peak perceptual difference. Note that the timing of the effects also coincides with the timing of the classifier results in terms of the maximum differences between perceptual and semantic categories (see Fig. 4). The ERP results thus mirror, qualitatively, the results of our previous multivariate analyses in terms of the timing of the maximum signal difference between categories. Again, these results suggest that perceptual aspects are coded in brain activity earlier than semantic aspects during visual processing, but semantic differences dominate the EEG signal earlier than perceptual ones during retrieval.

## 3. Discussion

When a memory is triggered by a reminder, how does its neural fingerprint unfold in time? While it is widely accepted that object recognition starts with low-level perceptual followed by high-level abstract processing (Carlson et al., 2013; Cichy et al., 2014; Lehky & Tanaka, 2016; Serre et al., 2007), much less is known about the mnemonic processing cascade. Here we demonstrate that the reconstruction of a visual memory does depend on a hierarchical stream too, but this mnemonic stream follows the reverse order relative to visual processing. Across three experiments, we found highly converging evidence in favour of such a reversal from behavioural reaction times and accuracy (Experiments 1 and 2), from multivariate classification analyses, and from univariate ERP analyses (Experiment 3).

The behavioural studies demonstrate that participants were significantly faster at detecting low-level perceptual differences than abstract, conceptual differences during a visual classification task, i.e. while an object was presented on the screen. Critically, however, when we asked participants to categorize the perceptual or semantic components of objects recalled from memory, the reverse effect was found: subjects required significantly less time to correctly retrieve semantic information about the object compared to perceptual details (see Fig. 2a and Fig. 2b). This reversal was corroborated by a significant interaction between the kind of feature (perceptual or semantic) and the kind of task (visual perception or memory recall task). Based on signal-detection models (Ashby, 2000; O’Connell, Dockree, & Kelly, 2012), the RT findings suggest that during memory reconstruction, the decision threshold to identify abstract information of a mnemonic representation is reached before a judgment about low-level information can be made. The response latency pattern therefore supports our central hypothesis that the temporal order of feature processing is reversed when retrieving a previously stored representation of an object, relative to its perception.

In addition to reaction times, the same reversal pattern was present in accuracy. Here, the accuracy profiles from Experiment 1 also allowed us to conduct a conditional probability analysis. Specifically, we were interested in whether access to semantic features and access to perceptual features are dependent on each other, and whether the direction of this mutual dependency would provide evidence for a processing hierarchy. Conditional probabilities revealed that when participants correctly retrieved perceptual information of a given object, they were highly likely to also make an accurate response about the semantic features of the same object, but not vice versa (see Fig. 2c). In other words, retrieving perceptual features required access to semantic features, but retrieving semantic features did not predict access to perceptual features to the same degree, as would be expected if the processing stream was hierarchically organized. These findings are consistent with an information-processing stream where access to perceptual details of a mnemonic representation depends on having completed the presumably earlier semantic stage, a finding consistent with hierarchical memory system models (Henson & Gagnepain, 2010).

The results from our third, EEG experiment fully support the conclusions drawn from the behavioural studies. We used temporally resolved multivariate decoding analyses to observe when in time, during object perception and object retrieval, the perceptual and semantic features of an object would be maximally decodable from a participant’s brain activity patterns. These analyses were carried out on a single trial level such that the fidelity peaks of the perceptual and semantic classifiers could be directly compared. When an object was visually presented during encoding, the maximum fidelity (*d* value) in classifying perceptual information (photograph vs. line drawings) occurred significantly earlier (approximately 100 ms) than the maximum for semantic information (animate vs. inanimate) (see Fig. 4a). This finding is consistent with a predominantly feed-forward processing stream as described previously (Carlson et al., 2013; Cichy et al., 2014; Clarke & Tyler, 2015; Lehky & Tanaka, 2016; Serre et al., 2007). Conversely, when we asked participants to reactivate an object’s representation from memory, peaks in classifying semantic information were found roughly 300ms before the peaks for perceptual categories (see Fig. 4b). This reversal in classifier fidelity was present on a trial-by-trial level but also when averaging peak latencies per participant (see Fig. 4c and Fig. 4d). Like in the behavioural experiments, a consistent reversal between perception and memory was supported by a significant interaction between the kind of feature (perceptual or semantic) and the type of task (perception vs. retrieval). Finally, we also found the same reversal pattern in the ERP peaks when comparing the maximum ERP difference between perceptual and semantic object classes. During object perception, the largest perceptual ERP cluster occurred ~100ms before the semantic ERP cluster, whereas during retrieval the perceptual cluster followed the semantic one with a lag of about 400ms (see Fig. 5). In summary, our two behavioural experiments, together with the decoding results and the ERP analyses, provide robust evidence for our main prediction that semantic features are prioritized over perceptual features during memory recall, opposite to the well-known forward stream of visual-perceptual processing. Follow-up studies will need to test whether this reversed stream is robust under different conditions, for example in tasks that vary the encoding demands to explicitly prioritize the encoding of perceptual over semantic aspects of an event.

In our studies, the behavioural data were acquired separately from the EEG data, in a setting that was optimized for measuring reaction times. Previous studies simultaneously measuring RTs and neural activity suggest that a meaningful relationship exists, on a single trial level, between the *d* values resulting from EEG classification and human behaviour. In line with signal detection models (Ashby, 2000; O’Connell et al., 2012), it has been argued that the distance between two or more categories in a neural representational space can serve as a decision boundary that guides behavioural categorization (Ritchie et al., 2015). For example, Carlson et al (Carlson, Ritchie, Kriegeskorte, Durvasula, & Ma, 2014) used fMRI-based activation patterns in late visual brain regions during an object recognition task, where participants had to make animacy judgements, similar to our semantic task. They found that the faster the reaction time on a given trial, the further away in neural space the object was represented relative to the boundary between semantic categories. Similarly, an MEG study (Ritchie et al., 2015) showed that the decision values during the time points of maximum decodability, derived in a way similar to our EEG study, were strongly correlated with reaction times for visual categorization. Both studies thus suggest that during object vision, single-trial decoding measures reflect a distance between categories in a neural space that directly translates into behaviour. Even though we did not obtain reaction times during the same trials that were used for EEG decoding, our findings indicate that this meaningful brain-behaviour relationship extends to mental object representations during memory reconstruction.

How does the reverse reconstruction hypothesis fit with existing knowledge about the neural pathways involved in memory reconstruction? It is generally accepted that during memory formation, information flows from domain-specific sensory modules via perirhinal and entorhinal cortices into the hippocampus. Recent evidence suggests that during visual processing, the coding of perceptual object information is preserved up to relatively late perirhinal processing stages (Martin, Douglas, Newsome, Man & Barense, 2018). The hippocampus is considered a domain-general structure (Howard Eichenbaum, 2004; Moscovitch, 2008; Staresina & Davachi, 2008) whose major role is the associative binding of the various elements that constitute an episode (Davachi, 2006; H. Eichenbaum, Yonelinas, & Ranganath, 2007; Squire, Stark, & Clark, 2004). The hippocampal code later allows a partial cue to trigger the reconstruction of these different elements from memory. This memory reconstruction process is thought to depend on back-projections from the hippocampus to neocortical areas, causing the reactivation of memory patterns in at least a subset of the areas that were involved in perceiving the original event. Such reactivation has consistently been reported in higher-order sensory regions related to processing of complex stimulus and task information (Johnson et al., 2009; Kuhl et al., 2011; Michelmann et al., 2016; Wimber et al., 2015), but also in relatively early sensory cortex (Bosch, Jehee, Fernandez, & Doeller, 2014; Waldhauser et al, 2016), suggesting that in principle higher- and lower-level information can be reconstructed from memory. Interestingly, however, recent evidence suggests that the semantic structure of complex naturalistic events is represented in brain activity patterns more consistently when participants reproduce the event narratives (movies) from memory, as opposed to watching the movies (Chen et al., 2017). Our work offers a neurobiologically plausible explanation for why higher-order meaningful information might be prioritized during retrieval. Within the medial temporal lobe, regions that are involved in the processing of objects and scenes are also activated when retrieving objects and scenes from memory, but with a delay relative to the actual perception of objects and scenes, consistent with a reversed information flow (Staresina, Cooper, & Henson, 2013). Intracranial EEG recordings have shown that connectivity between the entorhinal cortex and the hippocampus changes directionality between encoding and retrieval (Fell et al., 2016), which could provide the functional basis for cortical reinstatement. Studies in rodents also indicate that the neural codes that represent certain spatial trajectories are often replayed in reverse order when the animal is awake and resting, suggesting a potential role in memory retrieval (Carr & Frank, 2012), and there is very recent work in humans pointing to reverse replay of spatial sequences during offline states (Kurth-Nelson, Economides, Dolan, & Dayan, 2016). Finally, previous work using MEG decoding suggests that it is mainly the later processing stages of the encoding stream that are reactivated during retrieval, consistent with a prioritization of higher-level information during retrieval (Kurth-Nelson et al., 2015). Our proposal of a reverse processing hierarchy is thus plausible based on functional anatomy and the existing literature, even though it has never been explicitly proposed or tested so far.

We regard our reverse reconstruction hypothesis as complementary to existing models that address the nature and timing of different retrieval processes, including the influential dual process model (for a review see Yonelinas, Aly, Wang, & Koen, 2010). Dual process models focus on recognition rather than recall tasks, and on the cognitive processes and operations required to access a stored memory rather than the reactivated features of a memory themselves. They assume that successful recognition of a previously stored stimulus can be based on a sense of familiarity, or on the additional recollection of contextual information associated with the stimulus during encoding, an influential idea in the memory field since the introspective analyses of William James (James, 1890). While the original model does not explicitly address the time course of these processes, it is now widely accepted, based on the EEG literature, that familiarity signals occur earlier than recollection signals. Familiarity signals can be detected in the EEG as early as 300ms after the onset of a recognition probe, while recollection-related activity typically begins to emerge after 500-600ms (Bridson, Fraser, Herron, & Wilding, 2006; Klimesch et al., 2001; Mecklinger, 2006; Rugg & Curran, 2007). In contrast to the above-mentioned studies, our studies probed memory via cued recall, where successful recall strongly depends on the recollection of associative information. Within this recollection process, we find that the semantic “gist” of a memory is accessed before perceptual details. This hierarchical progression from an early global semantic (i.e., familiarity-like) signal to more fine-grained recollection might thus be a fundamental principle of retrieval that is shared between recall and recognition memory.

Beyond specific models of declarative memory, there are also interesting parallels between our findings and visual learning phenomena like the Eureka effect (Ahissar & Hochstein, 1997). The general idea that perception is shaped by stored representations has been proposed over a century ago by von Helmholtz (Helmholtz, 1924). A wealth of findings now support the idea that previous exposures to a stimulus can exert a strong top-down influence on its subsequent perception (for a review; Aggelopoulos, 2015). Reminiscent of our present findings, Ahissar and Hochstein (2004) suggest that such visual learning is a top-down process that progresses from high-level to low-level visual areas with increasing practice. Specifically, they argue that improvements in visual discrimination tasks (e.g. identifying a tilted line among distractors) are guided by high-level information (e.g. “the gist of the scene”) during earlier stages of learning, and increasingly more by low-level information (e.g. line orientations or colours) at later stages. Our findings indicate that during the reactivation of an object’s stored representation, its high-level features are retrieved more rapidly than its low-level components. Abstract information might thus be reactivated more easily and during earlier stages of visual learning, and thus have a stronger driving influence on performance than more detailed information. Even though speculative at the moment, our reverse reconstruction framework might thus have explanatory value for findings in related fields of learning and memory.

How our brain brings back to mind past events, and enriches our mental life with vivid images or sounds or scents beyond the current external stimulation, is still a fascinating and poorly understood phenomenon. Our present results suggest that memories, once they are triggered by a reminder, unfold in a systematic and hierarchical way, and that the mnemonic processing hierarchy is reversed with respect to the major visual processing hierarchy. We hope that these findings can inspire more dynamic frameworks of memory retrieval that explicitly acknowledge the reconstructive nature of the process, rather than simply conceptualizing memories as reactivated snapshots of past events. Such models will help us understand the heuristics and systematic biases that are inherent in our memories and memory-guided behaviours.

## 4. Methods

### 4.1. Participants

A total of 49 volunteers (39 female; mean age 20.02 +/- 1.55 years old) took part in behavioural Experiment 1. Twenty-six of them (19 female; mean age 20.62 +/- 1.62 years old) participated in the memory reaction time task. Five out of these 26 participants were not included in the final analysis due to poor memory performance (<66% general accuracy) compared with the rest of the group (*t*_24_ = 6.65, *p* < 0.01). Another group of 23 participants (20 female; mean age 19.35 ± 1.11 years) volunteered to participate in the visual reaction time task. In a second behavioural experiment (Experiment 2), 48 participants were recruited (42 female; mean age 19.25 +/- 0.91 years). Twenty-four of them performed the memory reaction time task and another group of 24 took part in the visual reaction time task. For the electrophysiological experiment we recruited a total of 24 volunteers (20 female; mean age 21.91 ± 4.68 years). Since the first 3 subjects we recorded performed a slightly different task during retrieval blocks (i.e., they were not asked to mentally visualise the object for 3 seconds, and they had to answer only one of the perceptual and semantic questions per trial), we did not include these participants in any of the retrieval analyses.

All participants reported being native or highly fluent English speakers, having normal (20/20) or corrected-to-normal vision, normal colour vision, and no history of neurological disorders. We received written informed consent from all participants before the beginning of the experiment. They were naïve as to the goals of the experiments, but were debriefed at the end. Participants were compensated for their time, receiving course credits or £6 per hour for participation in the behavioural task, or a total of £20 for participation in the electrophysiological experiment. The University of Birmingham’s Science, Technology, Engineering and Mathematics Ethical Review Committee approved all experiments.

### 4.2. Stimuli

In total, 128 pictures of unique everyday objects and common animals were used in the main experiment, and a further 16 were used for practice purposes. Out of these, 96 were selected from the BOSS database (Brodeur, Dionne-Dostie, Montreuil, & Lepage, 2010), and the remaining images were obtained from online royalty-free databases. All original images were pictures in colour on a white background. To produce two different semantic object categories, half of the objects were chosen to be animate while the other half was inanimate. Within the category of inanimate objects, we selected the same amount of electronic devices, clothes, fruits and vegetables (16 each). The animate category was composed of an equivalent number of mammals, birds, insects and marine animals (16 each). With the objective of creating two levels of perceptual manipulation, a freehand line drawing of each image was created using the free and open source GNU image manipulation software (www.gimp.org). Hence a total of 128 freehand drawings of the respective 128 pictures of everyday objects were created. Each drawing was composed of a white background and black lines to generate a schematic outline of each stimulus. For each subject, half of the objects were pseudo-randomly chose to be presented as photographs, and half of them as drawings, with the restriction that the two perceptual categories were equally distributed across (i.e. orthogonal with respect to) the animate and inanimate object categories. All photographs and line drawings were presented at the centre of the screen with a rescaled size of 500 × 500 pixels. For the memory reaction time task and the EEG experiment, 128 action verbs were selected that served as associative cues. Experiment 2 also used colour background scenes of indoor and outdoor spaces (900 × 1600 pixels) that were obtained from online royalty-free databases, which are irrelevant for the present purpose.

### 4.3. Procedure

#### 4.3.1. Behavioural experiments

##### 4.3.1.1. Experiment 1

###### Visual reaction time task

Before the start of the experiment, participants were given oral instructions and completed a training block of 4 trials to become familiar with the task. The main perceptual task consisted of 4 blocks of 32 trials each (Fig.1b). All trials started with a jittered fixation cross (500 to 1500ms) that was followed by a question screen. On each trial, the question could either be a perceptual question asking the participant to decide as quickly as possible whether the upcoming object is shown as a colour photograph or as a line drawing; or a semantic question asking whether the upcoming object represents an animate or inanimate object. Two possible response options were displayed at the two opposite sides of the screen (right or left). The options for “animate” and “photograph” were always located on the right side to keep the response mapping easy. The question screen was displayed for 3 seconds, and an object was then added at the centre of the screen. In Experiment 2, this object was overlaid onto a background that filled large parts of the screen. Participants were asked to categorize the object in line with the question as fast as they could as soon as the object appeared on the screen, by pressing the left or right arrow on the keyboard. Reaction times (RTs) were measured to test if participants were faster at making perceptual compared to semantic decisions.

All pictures were presented until the participant made a response but for a maximum of 10 sec, after which the next trial started. Feedback about participants’ performance was presented at the end of each experimental block. There were 256 trials overall, with each object being presented twice across the experiment, once together with a perceptual and once with a semantic question. Repetitions of the same object were separated by a minimum distance of 2 intervening trials. In each block, we asked the semantic question first for half of the objects, and the perceptual question first for the other half.

The final reaction time analyses only included trials with correct responses, and excluded all trials with an RT that exceeded the average over subjects by +- 2.5 standard deviations (SDs).

###### Memory reaction time tasks

The memory version was kept very similar to the visual reaction time task, but we now measured RTs for objects that were reconstructed from memory rather than being presented on the screen, and we thus had to introduce a learning phase first. At the beginning of the session, all participants received instructions and performed two short practice blocks. Each of the overall 16 experimental blocks consisted of an associative learning phase (8 word-object associations) and a retrieval phase (16 trials, testing each object twice, once with a perceptual and once with a semantic question). The associative learning and the retrieval test were separated by a distractor task. During the learning phase (Fig. 1c), each trial started with a jittered fixation cross (between 500 and 1500ms) that was followed by a unique action verb displayed on the screen (1500ms). After presentation of another fixation cross (between 500 and 1500ms), a picture of an object was presented on the centre of the screen for a minimum of 2 and a maximum of 10 seconds. Participants were asked to come up with a vivid mental image that involved the object and the action verb presented in the current trial. They were instructed to press a key (up arrow on the keyboard) as soon as they had a clear association in mind; this button press initiated the onset of the next trial. Participants were made aware during the initial practice that they would later be asked about the object’s perceptual properties as well as its meaning, and should thus pay attention to details including colour and shape. Within a participant, each semantic category and sub-category (electronic devices, clothes, fruits, vegetables, mammals, birds, insects, and marine animals) was presented equally often at each type of perceptual level (i.e. as a photograph or as a line drawing). The assignment of action verbs to objects for associative learning was random, and the occurrence of the semantic and perceptual object categories was equally distributed over the first and the second half of the experiment in order to avoid random sequences with overly strong clustering.

After each learning phase, participants performed a distractor task where they were asked to classify a random number (between 1 and 99) on the screen as odd or even. The task was self-paced and they were instructed to accomplish as many trials as they could in 45 seconds. At the end of the distractor task, they received feedback about their accuracy (i.e., how many trials they performed correctly in this block).

The retrieval phase (Fig. 1c) started following the distractor task. Each trial began with a jittered fixation cross (between 500 and 1500ms), followed by a question screen asking either about the semantic (animate vs. inanimate) or perceptual (photograph vs. line drawing) features for the upcoming trial, just like in the visual perception version of the task. The question screen was displayed for 3 seconds by itself, and then one of the verbs presented in the directly preceding learning phase appeared above the two responses. We asked participants to bring back to mind the object that had been associated with this word and to answer the question as fast as possible by selecting the correct response alternative (left or right keyboard press). If they were unable to retrieve the object, participants were asked to press the down arrow. The next trial began as soon as an answer was selected. At the end of each retrieval block, a feedback screen showing the percentage of accurate responses was displayed.

Throughout the retrieval test, we probed memory for all word-object associations learned in the immediately preceding encoding phase in pseudorandom order. Each word-object association was tested twice, once together with a semantic and once with a perceptual question, with a minimum distance of 2 intervening trials. In addition, we controlled that the first question for half of the associations was semantic, and perceptual for the other half. Like in the visual RT task, the response options for “animate” and “photograph” responses were always located on the right side of the screen. In total, including instructions, a practice block and the 16 learning-distractor-retrieval blocks, the experiment took approximately 60 minutes.

For RT analyses we only used correct trials, and excluded all trials with an RT that exceeded the average over subjects by +- 2.5 SDs.

##### 4.3.1.1. Experiment 2

Experiment 2 was very similar in design and procedures to Experiment 1, and we therefore only describe the differences between the two experiments in the following.

###### Visual reaction time task

The second experiment started with a familiarisation phase where all objects were presented sequentially. In each trial of this phase, a jittered fixation cross (between 500 and 1500 ms) was followed by one screen that showed the photograph and line drawing version of one object simultaneously, next to each other. During the presentation of this screen (2.5 sec) participants were asked to overtly name the object. After a jittered fixation cross (between 500 and 1500 ms), the name of the object was presented.

After this familiarisation phase, the experiment followed the same procedures as the visual reaction time task in Experiment 1 except for the following changes. Objects were overlaid onto a coloured background scene (1600 × 900 pixels). Also, each object (286 x 286 pixels) was probed only once, either together with a perceptual question, a semantic question (like above), or a contextual question asking whether the background scene was indoor or outdoor. For the current purpose we only describe the RTs to object-related questions in the Results section. Another minor difference to Experiment 1 was that in this version of the task, the question screen was displayed for 4sec, and the two options to answer during stimulus presentation were removed from the screen as soon as the object/reminder appeared.

###### Memory reaction time task

The memory reaction time task in Experiment 2 also included, during the associative learning phase, a background scene (1600 × 900 pixels) that was shown on the screen behind each object (286 × 286 pixels), and participants were asked to remember the word-background-object combination. In this version of the task, each word-object association was tested only once, together with either a perceptual question about the object, a semantic question about the object, or a contextual question regarding the background scene (indoor or outdoor). Therefore, one third of the objects were tested with a semantic question, one third with a perceptual question, and one third with a contextual question. Again, context was not further taken into account in the present analyses.

#### 4.3.2. EEG experiment (Experiment 3)

Following the EEG set-up, instructions were given to participants and two blocks of practice were completed. The task procedure of the EEG experiment was similar to the memory task in Experiments 1 and 2 except for the retrieval phase (Fig. 3a). Each block started with a learning phase where participants created associations between overall 8 action verbs and objects. After a 40 sec distractor task, participants’ memory for these associations was tested in a cued recall test. In total, the experiment was composed of 16 blocks of 8 associations each.

Each trial of the retrieval test started with a jittered fixation cross (500-1500ms), followed by the presentation of one of the action verbs presented during the learning phase as a reminder. Participants were asked to visualize the object associated with this action verb as vividly and in as much detail as possible while the cue was on the screen. To capture the moment of retrieval, participants were asked to press the up-arrow key as soon as they had the object back in mind; or the down-arrow if they could not remember the object. This reminder was presented on the screen for a minimum of 2 sec and until a response was made (maximum 7 sec). Immediately afterwards, a blank square with the same size as the original image was displayed for 3 sec. During this time, participants were asked to “mentally visualize the originally associated object on the blank square space”. After a short interval where only the fixation cross was present (500-1500ms), a question screen was displayed for 10 seconds or until participant response asking about perceptual (photograph vs. line drawing) or semantic (animate vs. inanimate) features of the retrieved representation, like in the behavioural tasks. However, in this case both types of questions were always asked on the same trial, and they were asked at the end of the trial rather than before the appearance of the reminder. The first question was semantic in half of the trials, and perceptual in the other half. Therefore, each retrieval phase consisted of 8 trials where we tested all verb-object associations learned in the same block in random order.

### 4.4. Data Collection (behavioural and EEG)

Behavioural response recording and stimulus presentation were performed using Psychophysics Toolbox Version 3 (Brainard, 1997) running under MATLAB 2014b (MathWorks). For response inputs we used a computer keyboard where directional arrows were selected as response buttons.

Electroencephalography (EEG) data was acquired using a BioSemi Active-Two amplifier with 128 sintered Ag/AgCl active electrodes. Through a second computer the signal was recorded at a 1024 Hz sampling rate by means of the ActiView recording software (BioSemi, Amsterdam, the Netherlands).

### 4.5. EEG Pre-processing

EEG data was pre-processed using the Fieldtrip toolbox (version from 3^rd^ August, 2017) for Matlab (Oostenveld, Fries, Maris, & Schoffelen, 2011). Data recorded during the associative learning phase was epoched into trials starting 500ms before stimulus onset and lasting until 1500ms after stimulus offset. The resulting signal was baseline corrected based on pre-stimulus signal (-500ms to onset). Retrieval epochs contained segments from 4000ms before until 500ms post-response. Since the post-response signal during retrieval will likely still contain task-relevant (i.e., object specific) information, we baseline-corrected the signal based on the whole trial. Both datasets were filtered using a low-pass filter at 100 Hz and a high-pass filter at 0.1 Hz. To reduce line noise at 50 Hz we band-stop filtered the signal between 48 and 52 Hz. The signal was then visually inspected and all epochs that contained coarse artefacts were removed. As a result, a minimum of 92 and a maximum of 124 trials remained per participant for the encoding phase, and a range between 80 and 120 trials per subject remained for retrieval. Independent component analysis was then used to remove eye-blink and horizontal eye movement artefacts; this was followed by an interpolation of noisy channels. Finally, all data was referenced to a common-average-reference (CAR).

### 4.6. Time resolved multivariate decoding

First, to further increase the signal to noise ratio for multivariate decoding, we smoothed our pre-processed EEG time courses using a Gaussian kernel with a full-width at half-maximum of 24ms. Time resolved decoding via linear discriminant analysis (LDA) using shrinkage regularization (Lemm, Blankertz, Dickhaus, & Müller, 2011) was then carried out using custom-written code in MATLAB 2014b (MathWorks). Two independent classifiers were applied to each given time window and each trial (see Fig. 3b): one to classify the perceptual category (photograph or line drawing) and one to classify the semantic category (animate or inanimate). In both decoding analyses, we used undersampling after artefact rejection (i.e. for the category with more trials we randomly selected the same number of trials as available in the smallest category). The pre-processed raw amplitudes on the 128 EEG channels, at a given time point, were used as features for the classifier. LDA classification was performed separately for each participant and time point using a leave-one-out cross-validation approach. This procedure resulted in a decision value (*d* value) for each trial and time point, where the sign indicates in which category the observation had been classified (e.g., - for photographs and + for line drawings in the perceptual classifier), and the value of *d* indicates the distance to the hyper-plane that divided the two categories (with the hyper-plane being 0). This distance to the hyper-plane provided us with a single trial time-resolved value that indicates how confident the classifier was at assigning a given object to a given category. In order to use the resulting *d* values for further analysis, the sign of the *d* values in in one category was inverted, resulting in d-values that always reflected correct classification if they had a positive value, and increasingly confident classification with increasingly higher values.

Our main intention was to identify the specific moment within a given trial at which each of the two classifiers showed the highest fidelity, and to then compare the temporal order of the perceptual and semantic peaks. We thus found the maximum positive *d* value in each trial and separately for the semantic and perceptual classifiers, with the important restriction that we only used peaks with a value exceeding the 95^th^ percentile of the classifier chance distribution (see section on bootstrapping below), such as to minimize the risk of including meaningless noise peaks. The resulting output from this approach allowed us to track and compare the temporal “emergence” of perceptual and semantic classification within each single-trial. In addition to this single-trial analysis, we also calculated the average *d* value peak latency for perceptual and semantic classification in each participant to compare the two average temporal distributions. Note, however, that many factors could obscure differences between semantic and perceptual peaks when using this average approach, including variance in processing speed across trials, e.g. for more or less difficult recalls. We therefore believe that the single trial values are more sensitive to differences in timing between the reactivated features.

### 4.7. Generating an empirical null distribution for the classifier

Previous work has shown that the true level of chance performance of a classifier can differ substantially from its theoretical chance level that is usually assumed to be 1/number of categories (Combrisson & Jerbi, 2015; Jamalabadi, Alizadeh, Schönauer, Leibold, & Gais, 2016; Kowalczyk & Chapelle, 2005). A known empirical null distribution of *d* values would allow us to determine a threshold for considering only those *d* value peaks as significant whose values are higher than the 95^th^ percentile of this null distribution. We generated such an empirical null distribution of *d* values by repeating our classifier analysis with randomly shuffled labels a number of times, and combined this with a bootstrapping approach, as detailed in the following.

As a first step, we generated a set of d-value outputs that were derived from carrying out the same decoding procedure as for the real data (including the leave-one-out cross-validation), but using category labels that were randomly shuffled at each repetition. This procedure was carried out independently per participant. On each repetition, before starting the time-resolved LDA, all trials were randomly divided into two categories with the constraint that each group contained a similar number of photographs and line drawings, and approximately the same amount of animate and inanimate objects (the difference in trial numbers was smaller than 8%). The output of one such repetition per participant was one d-value per trial and time-point, just as in the real analysis. This procedure was conducted 50 times per participant for object perception (encoding) and retrieval, respectively, with a new random trial split and random label assignment on each repetition. For each participant we thus had a total of 51 classification outputs, one using the real labels, and 50 using the randomly shuffled labels.

Second, we also used the shuffled label outputs in order to generate an empirical Z-score distribution for our single-trial analyses. Our main statistic of interest with respect to the EEG data was a Wilcoxon signed rank test comparing the order of the perceptual and semantic classifier peaks on each single trial. This analysis was based on all available single trials accumulated across participants, and thus resulted in a high number of degrees of freedom, with a possibly exaggerated likelihood of finding a significant Z-score. We therefore tested our real data against an empirical Z-score distribution obtained from a series of bootstrapping analyses that were based on the same data and simulated the same number of degrees of freedom. For each participant’ trial, we took the outputs from two different classifiers randomly selected from a sample of 52 classifiers (i.e., 50 with shuffled labels, one real perceptual, and one real semantic). That is, we created two arbitrary conditions per trial to make a pairwise comparison (emulating our perceptual vs. semantic conditions). There was a 50:1 chance that the “pseudo-semantic” classifier contained the output of the real semantic classifier, and likewise a 50:1 chance that the “pseudo-perceptual” classifier contained the d-values from the real perceptual classifier. Next, we choose for each type of condition the highest *d* value per trial in the accurate direction and in a given time window, using the same constraints as for the real classifier outputs. This provided us with one peak per condition (two) for every trial. To equate the number of degrees of freedom with our contrast of interest, we randomly selected the same number of pairs as available in the real analysis. Finally, a Wilcoxon signed rank test was used to compare the temporal distance of the *d* value peaks between the two conditions, and the corresponding Z-value was registered, again mirroring the analysis carried out on the real data. This approach was repeated with replacement for a total of 10000 times, generating an empirical distribution of Z-values under the null hypothesis that there is no meaningful information about an object’s category in the EEG data.

Thirdly, to estimate our classification chance distribution for the random-effects (i.e., trial-averaged) peak analyses, we used the 51 classification outputs from all participants in a bootstrapping procedure (Stelzer, Chen, & Turner, 2013). On each of the bootstrapped repetitions, we randomly selected one of the 51 classification outputs (50 from shuffled labels classifiers and one from a real labels classifier) per participant, and calculated the *d* value group average based on this random selection for each given time point. This procedure was repeated with replacement 10000 times. To generate different distributions for the perceptual and semantic classifiers, we run this bootstrapping approach two times: once where the real labels output from each subject came from the semantic classifier, and once where the real *d*-values came from the perceptual classifier.

### 4.8 Univariate event-related potential (ERP) analysis

A series of cluster-based permutation tests (Monte Carlo, 2000 repetitions, clusters with a minimum of 2 neighbouring channels within the FieldTrip software) was carried out in order to test for differences in ERPs between the two perceptual (photograph vs. line drawing) and the two semantic (animate vs. inanimate) categories, controlling for multiple comparisons across time and electrodes. First, we contrasted ERPs during object presentation in the encoding phase in the time interval from stimulus onset until 500ms post-stimulus. We then carried out the same type of perceptual and semantic ERP contrasts during retrieval, in this case aligning all trials to the time of the button press. We used the full time window from 3000ms before until 100ms after the button press, but we further subdivided this time window into smaller epochs of 300ms to run a series of T-tests, again using cluster statistics to correct for multiple comparisons across time and electrodes. We were mainly interested in the temporal order of the ERP peaks that differentiated between perceptual and semantic classes during encoding and retrieval. These peaks are based on statistically meaningful clusters as described above, but we conducted no further statistical comparisons between the average perceptual and semantic ERP peaks.

### 4.9 Data and code availability statement

The data and the custom code that support the findings of this study are available from the corresponding author upon reasonable request.

## Acknowledgments

We thank Alexandru-Andrei Moise, Emma Sutton, Thomas Faherty, Laura De Herde and James Lloyd-Cox for helping with data collection, and Rodika Sokoliuk for her useful technical support. This work was supported by an European Research Council Starting Grant (ERC-2016-STG-715714) and a scholarship from the Midlands Integrative Biosciences Training Partnership (MIBTP), which is a Biotechnology and Biological Sciences Research Council (BBSRC) funded doctoral training programme.

## Author contributions

J.L.D. and M.W. designed the experiments. J.L.D. conducted the experiment. J.L.D., M.S.T. and C.K. analysed the data. All authors contributed to the analysis approach and to data interpretation. J.L.D. and M.W. wrote the first version of the manuscript and all authors contribute in reviewing and editing.

